# Modular Thermal Control of Protein Dimerization

**DOI:** 10.1101/694448

**Authors:** Dan I. Piraner, Yan Wu, Mikhail G. Shapiro

## Abstract

Protein-protein interactions and protein localization are essential mechanisms of cellular signal transduction. The ability to externally control such interactions using chemical and optogenetic methods has facilitated biological research and provided components for the engineering of cell-based therapies and materials. However, chemical and optical methods are limited in their ability to provide spatiotemporal specificity in light-scattering tissues. To overcome these limitations, we present “thermomers,” modular protein dimerization domains controlled with temperature – a form of energy that can be delivered to cells both globally and locally in a wide variety of in vitro and in vivo contexts. Thermomers are based on a sharply thermolabile coiled-coil protein, which we engineered to heterodimerize at a tunable transition temperature within the biocompatible range of 37–42 °C. When fused to other proteins, thermomers can reversibly control their association, as demonstrated via membrane localization in mammalian cells. This technology enables remote control of intracellular protein-protein interactions with a form of energy that can be delivered with spatiotemporal precision in a wide range of biological, therapeutic and living material scenarios.

## INTRODUCTION

The study and engineering of cellular function within the context of complex tissues and synthetic biomaterials necessitates the development of methods to enable external control of cellular signaling with high spatiotemporal specificity and deep tissue penetration^1^. Conditional proteinprotein interactions are among the most widespread and versatile modalities employed by cells to regulate molecular signaling pathways^2^. Consequently, engineered protein associations have served as important tools in studying cellular signaling and constructing synthetic cellular devices. In particular, widely used chemically-inducible dimerization domains such as FKBP/FRB^3^ have enabled a vast array of applications ranging from the basic study of protein function^4^ to the engineering of exogenously-gated chimeric antigen receptors for cellular immunotherapy^5^. Chemical dimerization is effective both in culture and in deep tissues based on the bioavailability of the chemical agent; however, this activity is poorly amenable to spatiotemporal regulation. In contrast, optically inducible dimerization domains such as Cry2/CIB1 offer exquisite spatial and temporal precision, enabling microscopic studies of processes such as the immune response^6^ and cell motility^7^, but are limited by the scattering of photons in deep tissue and other complex media.

Temperature offers an alternative mechanism for controlling biological signaling with several advantages over chemicals and light. Temperature can be applied to biological samples globally using simple heat sources or electromagnetic radiation, and can be targeted locally deep within scattering media using technologies such as focused ultrasound, providing spatial and temporal resolution on the order of millimeters and seconds, respectively^1,8^. Previous work on thermal control of cellular signaling has focused on temperature-actuated transcription and translation, taking advantage of endogenous heat shock promoters^9,10^, temperature-dependent RNA elements^11^, or heterologously expressed protein-based transcriptional bioswitches^12^. In addition, temperature-sensitive variants of individual proteins have been used to study the function of these proteins in model systems^13,14^. While these pioneering approaches enable thermal control of specific aspects of cellular function, they lack the modularity of chemical and optical dimerizers.

Here we introduce a modular approach to controlling protein dimerization with temperature. Starting with a homodimeric temperature-dependent coiled-coil transcription factor from bacteria, we engineer a pair of heterodimeric protein association domains with sharp, tunable thermal unbinding. We demonstrate the ability of these “thermomers” to dynamically control protein localization in living mammalian cells. The resulting technology has the potential to provide versatile thermal control of protein-protein interactions in a variety of cell types, complementing the existing set of chemical and optical tools.

## RESULTS

As a starting point for the design of thermomers, we used TlpA, a temperature-sensitive transcriptional repressor from *Salmonella typhimurium* with a relatively simple architecture and well-characterized thermal behavior^15^. While no atomic-resolution structure of this protein exists, biochemical and bioinformatic studies have indicated that TlpA consists of an N-terminal DNA-binding domain and a C-terminal coiled-coil domain, the latter of which causes the protein to homodimerize in a temperature-dependent manner. As a dimer, TlpA binds to its cognate DNA operator and prevents transcription of the downstream gene. The coiled-coil domain of TlpA undimerizes and uncoils above a temperature of ~42°C, causing unbinding from the operator and transcriptional de-repression. This sharply cooperative transition happens over less than 3 °C, as defined by 10% to 90% activation. We previously demonstrated that the thermal set-point of TlpA could be tuned through directed evolution without compromising cooperativity, and used its transcription factor activity to spatiotemporally control the function of engineered bacteria with ultrasound hyperthermia^12^.

To turn TlpA into a modular protein-protein dimerization system, we first needed to convert it from a homodimer to a heterodimer. Most applications of inducible dimerization systems require the interacting modules to be heterodimeric to enforce selective binding between two desired molecular partners^16^. To redesign the wild type TlpA into a pair of heterodimeric coiled-coil species (**Fig. 1a**), we used rational mutagenesis guided by bioinformatic prediction of the TlpA dimerization domain structure. Coiled-coil domains typically consist of repetitive seven-amino acid residue sequences known as heptad repeats^17^. We used a published annotation of the TlpA sequence^18^, cross-referenced against a computational annotation from the COILS^19^ prediction server, to establish the register of heptad repeats within the TlpA primary sequence. We then introduced charge-complementary pairs of residues^20^ predicted either at conventional g-to-e’ contacts or at alternative g-to-d’ interfaces to disfavor homodimerization and favor heterodimerization (**Fig. 1, b-c**). The latter architecture occurs when large ionic sidechains at the core peripherally expose their charged termini, as has been described for the Fos-Jun coiled coil interaction^21^. To maintain the highly switch-like thermal dissociation behavior of TlpA, we reasoned that the least perturbative positions for mutagenesis would be at existing interfacial ionic interaction sites in the wild-type protein, which are present due to the C_2_ symmetry of the parallel coiled-coil structure. We mutagenized all such positions one by one, replacing cationic residues with glutamate and anionic side chains with arginine or lysine. We expressed the resulting coiled-coil domains in *E. coli*, purified the proteins via affinity chromatography, and assayed their helical content over a relevant thermal range using circular dichroism spectroscopy (**Supplementary Fig. S1**). From this initial screen we obtained a pair of charge-complemented mutants, dubbed TlpA-G_1_A (E180R) and TlpA-G_1_B (R179E), that demonstrated a sharp, sigmoidal thermal response profile with a notable upshift in the temperature threshold for an equimolar mixture of the mutant pair relative to pure solutions of either species (**Fig. 1d**).

**Figure 1:**
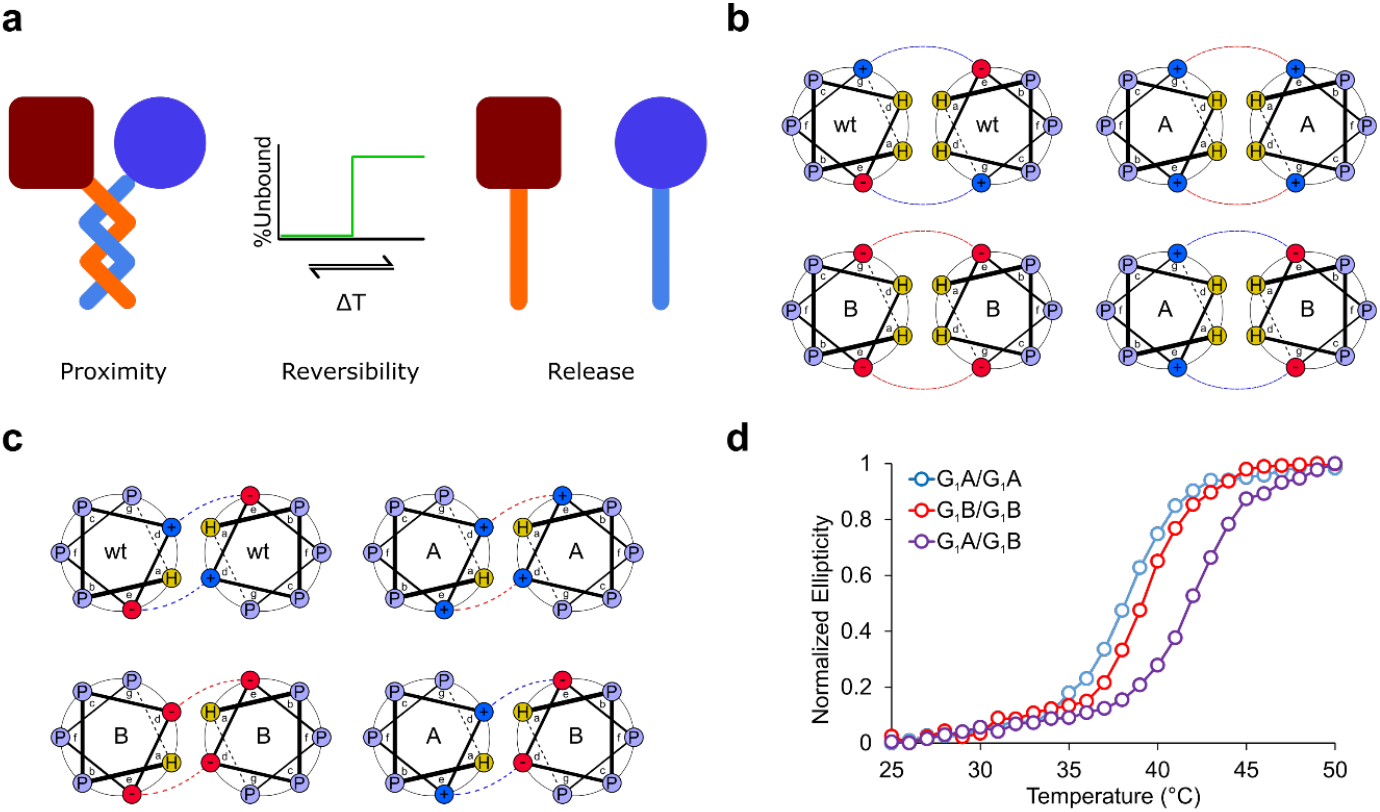
Engineering heterodimeric TlpA variants via charge-charge complementation. **a**) Illustration of TlpA-based thermomer system. Heterodimeric coiled-coil domains enable reversible association and dissociation of fusion partners as a sharp function of temperature. **b**) Diagram of heterodimeric coiled-coil design based on the introduction or modification of electrostatic contacts at the e-to-g’ interface between adjacent α-helices. P, H, + and — denote polar, hydrophobic and charged residues, respectively. WT denotes the wild-type protein, while A and B denote engineered mutants. **c**) Diagram of predicted electrostatic contacts along the TlpA interface occurring in a nonconventional e-to-d’ configuration. **d**) Normalized ellipticity of purified TlpA coiled-coil domain variants in isolation or as an equimolar mixture, measured at the 222 nm peak for α-helical spectra as a function of temperature. Data shown normalized from 0 to 1 on a per-sample basis.

To validate the *in cellulo* functionality of the TlpA-G_1_A and G_1_B mutants, we utilized the ability of TlpA to modulate the expression of a fluorescent reporter gene in *E. coli*^12^. We constructed a temperature-inducible circuit containing two separate copies of the TlpA gene, with TlpA operators upstream of a green fluorescent protein (GFP) and a red fluorescent protein (RFP). To compare the repression efficiency of the G_1_A/G_1_A and G_1_B/G_1_B homodimers to that of the G_1_A/G_1_B heterodimer, we generated circuit variants containing two copies of TlpA-G_1_A, two copies of TlpA-G_1_B, or one copy of each TlpA variant (**Fig. 2, a-b**). The thermal gene expression profiles of GFP showed all three circuits to produce a highly switch-like cooperative activation. However, the two homodimeric constructs had a clear downshift in their transition temperature compared to the heterodimeric construct containing both TlpA variants (**Fig. 2c**), confirming a stabilized heterotypic association between the two coiled-coil strands. The RFP output displayed similar activation profiles (**Supplementary Fig. S2**). Swapping the positions of the two TlpA copies within the vector did not significantly influence the expression profile, controlling for inadvertent stoichiometric effects (**Supplementary Fig. S3**).

**Figure 2:**
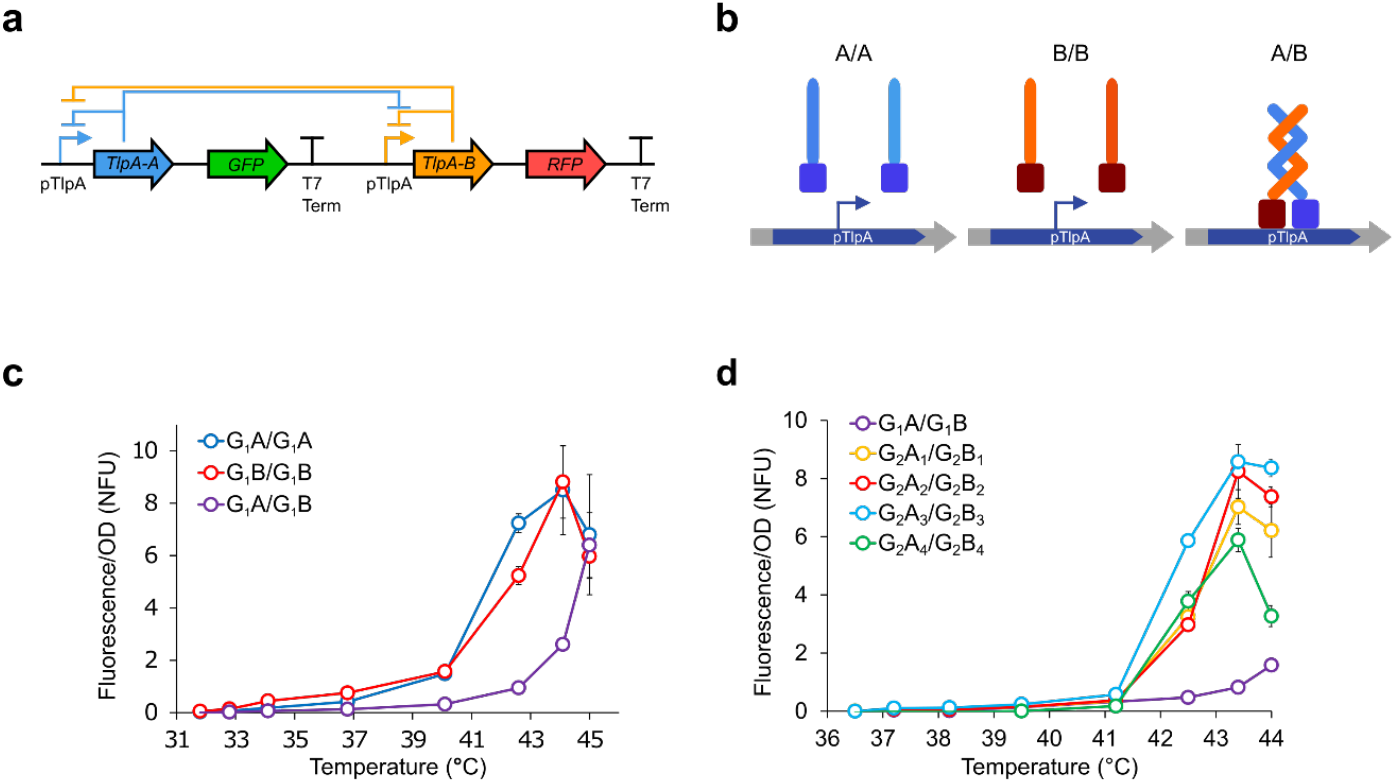
Evaluation of TlpA heterodimerization via reconstitution of promoter repression in bacteria. **a**) Diagram of bacterial genetic circuit containing two TlpA genes, each of which can encode one of the engineered variants. Bacteria harboring the G_1_A/G_1_A and G_1_B/G_1_B circuits can only produce the homodimeric coils, whereas the G_1_A/G_1_B circuits can produce either the homodimers or heterodimeric coils. **b**) Illustration of operator binding for engineered heterodimeric construct; only cells expressing both partners of the heterodimeric TlpA variant can repress the reporter gene above a certain temperature. **c**) OD-normalized fluorescence expression profile of *E. coli* harboring the plasmid constructs shown in **a**, with the single mutant G_1_A and G_1_B variants in the A and B positions, as a function of temperature (n = 3). Error bars represent ± s.e.m. NFU represents normalized fluorescent units. **d)** OD-normalized temperature-dependent fluorescence expression profiles of *E. coli* harboring plasmids bearing the four possible double mutant combinations of the G_1_ variants and two additional candidate single mutations, all in heterodimeric pairings (n = 3). Error bars represent ± s.e.m.

While our first-generation variants demonstrated the ability to engineer a heterodimerization preference, we noted that the G_1_A/G_1_A and G_1_B/G_1_B circuit constructs still had a transition setpoint above 37 °C, indicating that these mutants retained the ability to homodimerize under typical mammalian homeostatic conditions. We therefore used our thermal GFP expression assay to evaluate a subset of additional rational mutant pairs selected from our original panel (**Supplementary Fig. S4**), and chose the two best-performing pairs of substitutions from this subset to combine with TlpA G_1_A and TlpA G_1_B in all possible permutations. This resulted in the second-generation coiled-coil pairs dubbed TlpA G_2-1_ – G_2-4_, each comprising a G_2_A_n_ and a G_2_B_n_ monomer (**Supplementary Table 1**). In our bacterial bioswitch assay, all the heterodimeric circuits combining G_2_A_n_ with its complementary G_2_B_n_ displayed switch-like activation of reporter fluorescence (**Fig. 2d, Supplementary Fig. S5**). In contrast, the homodimeric constructs containing two copies of G_2_A_n_ or G_2_B_n_ were unable to propagate in a stable manner, consistently displaying deletions in the TlpA promoter or the fluorescent reporter gene, even when grown at 30 °C in recombination-deficient *E. coli*. We interpreted this as evidence that the second-generation variants are unable to form homodimeric interactions at the concentrations defined by our circuits, resulting in constitutive expression from the TlpA promoter and an untenable metabolic burden to the host cell^22,23^.

To confirm the dimerization preference of our engineered coiled-coils, we designed a biochemical assay based on covalent crosslinking and size-based gel separation. TlpA dimers can be crosslinked via CuCl_2_-catalyzed oxidation of the protein’s single cysteine residue^24^. To distinguish hetero-from homodimerization, we truncated one of the two TlpA sequences by removing its DNA binding domain, thereby altering its electrophoretic mobility on a polyacrylamide gel without perturbing its ability to dimerize (**Fig. 3a**). HA tags were added at the C-termini of both proteins to facilitate specific detection via Western blotting. We expressed the resulting pairs of truncated and full-length TlpA variants in *E. coli*. To validate this assay, we expressed a pair of wild-type TlpA coils and crosslinked them at 37 °C, or after a thermal elevation to 45 °C, or after return to 37 °C. Three bands corresponding to the expected mixture of the two types of homodimers and one type of heterodimer were visible after crosslinking at 37 °C, while crosslinking at the higher temperature resulted in a preponderance of monomers, which could be re-annealed by bringing the temperature back down to 37 °C (**Fig. 3b**).

**Figure 3:**
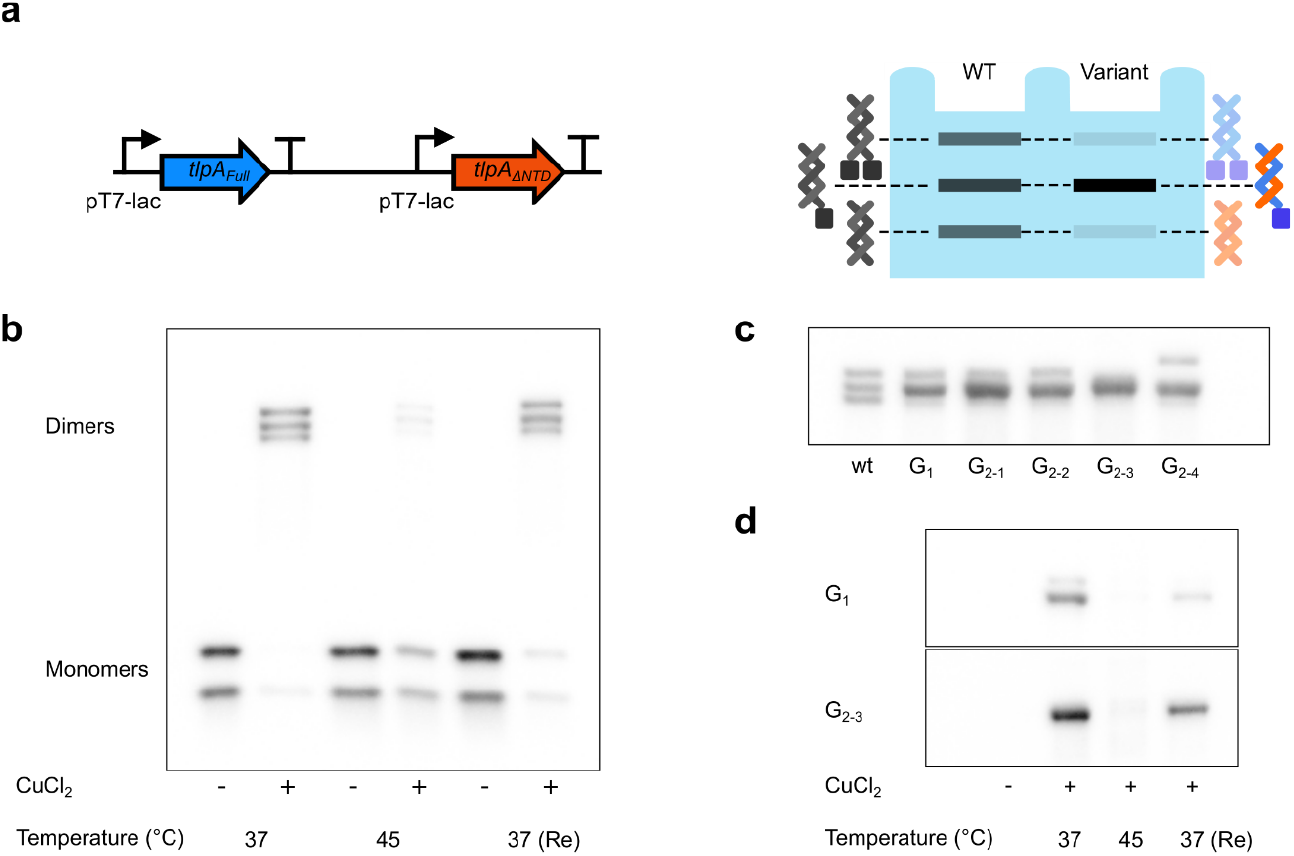
Validation of TlpA heterodimerization by electrophoresis. **a)** Left: diagram of genetic construct for simultaneous coexpression of engineered heterodimers of TlpA for biochemical assay. One open reading frame expresses the full length TlpA protein whereas the other ORF produces a truncated version missing its predicted N-terminal DNA-binding domain (ΔNTD). Each position can be occupied by any variant of TlpA, including the wild type protein and engineered mutants. Right: Diagram of the possible SDS-PAGE bands resulting from covalent dimeric crosslinking of the TlpA products expressed from this construct. The example in the left lane corresponds to the wildtype homodimeric TlpA. The example in the right lane corresponds to a pair of heterodimeric TlpA variants. **b)** Western blot of CuCl_2_-catalyzed crosslinking reaction of wild-type TlpA in *E. coli* lysate followed by SDS-PAGE. Crosslinking was performed at 37 °C, at 45 °C, or at 37 °C to assess reannealing (Re) following a 10-minute incubation at 45 °C. Each condition is compared to a non-crosslinked control. The bottom bands on the gel show uncrosslinked monomers. **c)** CuCl_2_ crosslinking, SDS-PAGE, and Western blot of the construct in **a** harboring wild type TlpA (wt), the first generation single mutant heterodimer (G_1_), or the G_2-n_ double mutant heterodimers. Crosslinking was performed at 37 °C. **d)** Thermal response of the G_1_ and G_2-3_ heterodimer constructs analyzed in the absence and presence of CuCl_2_ at 37 °C, 45 °C, or with 37 °C reannealing after a 10-minute incubation at 45 °C.

Substituting the wild type coiled-coils with our first-generation hetrodimerizing mutants resulted in preferential accumulation of the TlpA heterodimer at 37 °C (**Fig. 3c**). Constructs containing the second-generation variants demonstrated further reduction in the intensity of the homodimer bands in favor of the intermediate molecular weight heterodimer, with TlpA G_2-3_ demonstrating the strongest heterodimeric enrichment. (**Fig. 3c, Supplementary Fig. S6**). The first- and second-generation heterodimers both showed reversible dissociation at 45°C (**Fig. 3d**). On the basis of these results, the TlpA G_2-3_ pair was chosen as the “thermomer” construct for further experiments.

After validating the temperature-dependent association of our engineered heterodimeric TlpA thermomers, we set out to demonstrate their ability to be fused with other proteins and confer controlled protein-protein association in mammalian cells. We designed a construct wherein one TlpA-G_2-3_ strand was N-terminally fused with the palmitoylation sequence of GAP43, thereby compartmentalizing it to the plasma membrane. The complementary TlpA-G_2-3_ strand was fused at the C-terminus to mScarlet-I (**Fig. 4a**), an RFP chosen for its robust fluorescence at elevated temperature (**Supplementary Fig. S7**). To make this system compatible with mammalian homeostatic conditions, we combined the TlpA-G_2-3_ heterodimerizing mutations with three previously described amino acid substitutions that lower the coiled-coil dissociation temperature to approximately 39 °C^12^.

**Figure 4:**
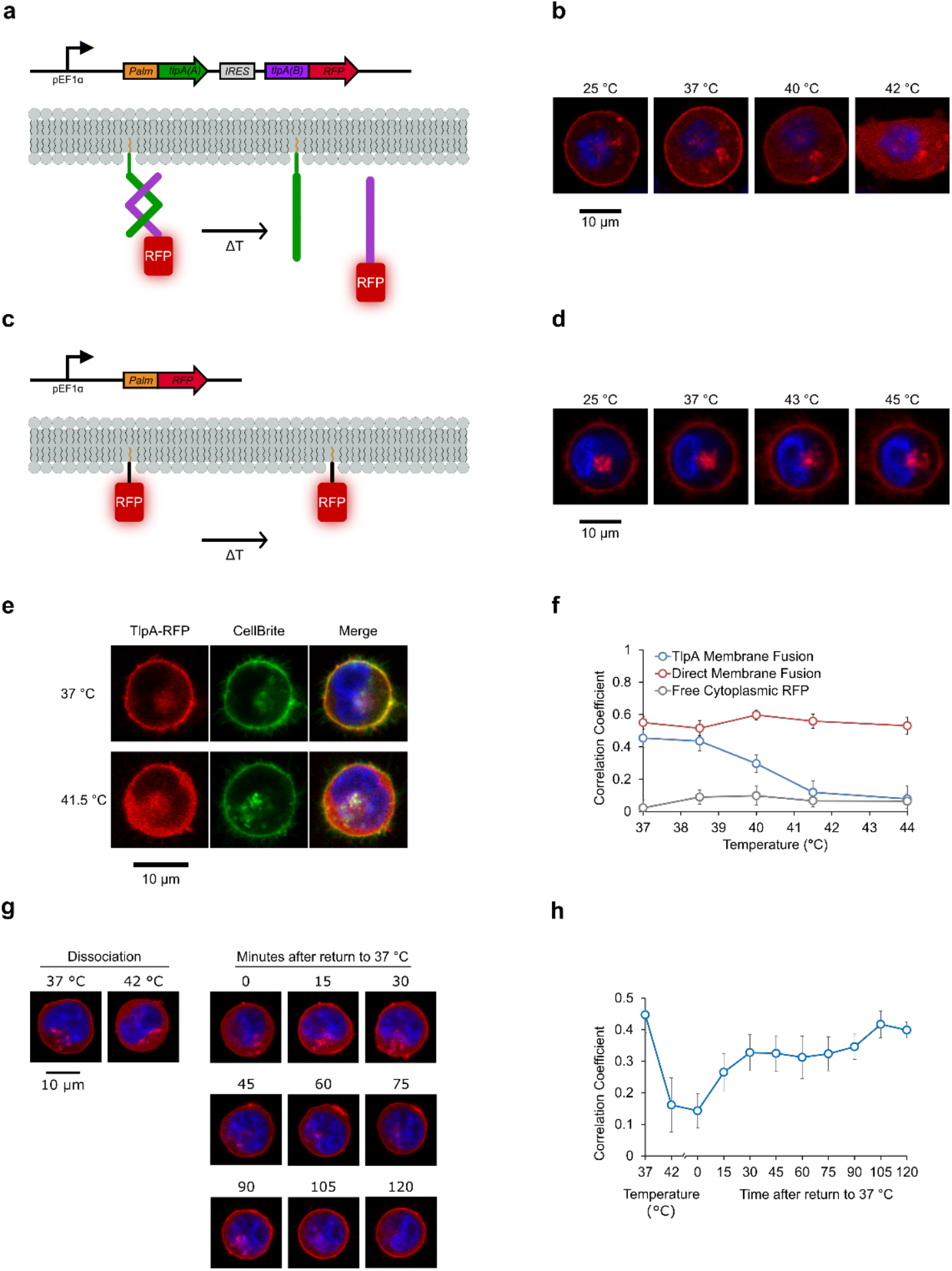
Membrane localization assay for TlpA activity in mammalian cells. **a**) Genetic construct (top) and schematic (bottom) for temperature-dependent localization of RFP at the plasma membrane. The TlpA_39_-G_2_A_3_ strand is fused to a GAP43 palmitoylation motif, leading to its tethering to the lipid membrane. The partner TlpA_39_-G_2_B_3_ strand is fused to RFP. At low temperature, the heterodimerization of these strands should lead to RFP localization at the membrane. Upon heating, the RFP-fused strand should dissociate from the membrane. **b**) Fluorescence images of a representative K562 cell transfected with the construct shown in **a**. Robust membrane localization of RFP fluorescence is observed at 25 °C and 37 °C. At 40 °C, RFP begins to dissociate from the membrane, and by 42 °C red fluorescence is distributed throughout the cytoplasm. The nucleus is stained in blue with Hoechst 33342 dye. **c**) Genetic construct (top) and experimental schematic (bottom) of a directly palmitoylated RFP control. **d**) Fluorescence images of representative K562 cell transfected with the construct shown in **c,** displaying robust membrane localization throughout the temperature range tested. **e**) Representative fluorescence images of a K562 cell with a lentivirally delivered TlpA-mediated RFP localization construct (shown in **a**) and the membrane-staining dye CellBrite Fix 488. **f**) Pixel colocalization of RFP with the CellBrite 488 dye as a function of temperature in K562 cells stably expressing directly palmitoylated RFP (shown in **b**, n = 8), the TlpA-mediated membrane localization construct (shown in **a**, n = 15), and free cytosolic RFP (n = 8). Error bars represent ± s.e.m. **g**) Time series of fluorescent images of a representative K562 thermal reporter cell line before heating, after heating to 42 °C, and after re-equilibration to 37 °C. RFP re-accumulation at the plasma membrane was tracked in 15 minute intervals. Pixel intensity was normalized to the maximal per-image value. **h)** Pixel colocalization of the TlpA-RFP signal with CellBrite 488 dye as a function of temperature (dissociation) followed by a time course of incubation at 37 °C reassociation; n = 12. Error bars represent ± s.e.m.

Using live cell confocal microscopy of transiently transfected K562 cells, at physiological temperature we observed strong localization of red fluorescence to the plasma membrane (**Fig. 4b** and **Supplementary Fig. S8**) and also to the Golgi apparatus (**Supplementary Fig. S9**). Increasing the cells’ temperature using resistive heating above a threshold of 40 °C resulted in the redistribution of membrane fluorescence into the cytosol. As a control for non-specific thermal dissociation, cells in which the RFP was directly palmitoylated showed no redistribution of membrane fluorescence within this temperature range (**Fig. 4, c-d**, and **Supplementary Fig. S10**). This confirms that RFP dissociation from the membrane is driven by a TlpA-mediated binding transition rather than disruption of membrane integrity.

To enable the use of the thermomer system with viral gene delivery and genomic integration, we generated a nonhomologous variant of the TlpA G_2_A_3_ strand (nhTlpA G_2_A_3_) in which all degenerate codons were mutagenized to synonymous triplets with minimal identity to the original sequence. This mutagenesis helps avoid template switching-mediated recombination during viral delivery of high-homology constructs^25^. The resulting open reading frame had 57.48% sequence identity to the parent sequence, with no more than 5 consecutive homologous nucleotides. Lentiviral delivery of a construct containing palmitoylated nhTlpA G_2_A_3_ and mScarlet-fused TlpA G_2_B_3_ resulted in robust temperature-induced dissociation of red fluorescence from the plasma membrane (**Supplementary Fig. S11**), similar to the results of transient transfection.

We used this virally-engineered K562 cell line to quantify the co-localization of RFP fluorescence intensity with signal from the plasma membrane, as delineated by CellBrite Fix 488 staining (**Fig. 4e**). Cells with thermomer-mediated RFP targeting to the membrane demonstrated co-localization with the dye at physiological temperature, followed by loss of pixel correlation above 40 °C (**Fig. 4f**). In contrast, control cells with directly palmitoylated RFP demonstrate robust colocalization with the membrane stain throughout the temperature range tested, while free cytoplasmic RFP showed no correlation with the CellBrite dye (**Fig. 4f**). We also used the TlpA reporter cell line to evaluate the reversibility of TlpA-mediated membrane localization after heating. Membrane localization was released by a 5-minute incubation at 42 °C. Upon cooling back to 37 °C, TlpA re-partitioned to the plasma membrane, indicating reversibility, albeit with slower kinetics than observed for dissociation (**Fig. 4, g-h**).

## DISCUSSION

Taken together, our results establish engineered heterodimeric TlpA coils as modular, tunable thermomers capable of conferring temperature-controlled protein-protein association and localization to genetically fused proteins. This technology complements the large existing repertoire of chemical and optical dimerizers used to control a wide range of protein andcellular functions^26^, as well as previous work on temperature-controlled transcription^12^. We anticipate that as external control of protein signaling becomes needed in more complex settings, such as cellular therapy^27^ and engineered living materials^28^, this will provide a role for temperature-based control modalities that offer spatiotemporal specificity and penetration depth beyond those afforded by systemic drug delivery and optical methods^1,8^.

We anticipate that the coiled-coil structure of TlpA will facilitate future use of TlpA-based thermomers to control a variety of protein functions. Coiled-coils have seen a diverse array of applications in biomaterials, affinity purification, assembly of bispecific or high avidity antibodies, and reconstitution of various split proteins or dimerization-dependent protein complexes^29,30^. Conditionally dimeric coiled-coil fusions have previously been used to reconstitute and control a split mutant of RNAse T1^31^ and to conditionally enact DNA binding by a bZIP domain upon photostimulation^32^.

To maximize its versatility as a tool for biology and medicine, future work on TlpA-based thermomers is needed to optimize construct size, modularity and kinetics. At 371 residues, TlpA is nearly four times larger than its drug-responsive counterpart, FKBP, which may hinder applications in gene delivery where sequence compactness is of premium importance. Truncation and mutagenesis may offer a strategy by which the size of the thermomer system could be minimized without sacrificing its sharp thermal switching response. In addition, elucidating the mechanism responsible for the unique thermal cooperativity of TlpA would assist in future rational engineering efforts to modify performance and generate additional, orthogonal thermomers. Furthermore, systematic characterization of potential genetic fusion sites and optimal linker sequences would facilitate the widespread adoption of TlpA as a thermal fusion moiety. In this study, we demonstrated that TlpA retains a cooperative thermal transition as an intact protein, as an isolated coiled-coil domain, and as a system with distinct fusion sites at the N and C-termini. However, additional engineering may be needed to accommodate fusion partners of different sizes and valency. Finally, it will be useful to study the factors governing thermal dissociation and re-association kinetics. In our experiments monitoring TlpA-RFP delocalization from the plasma membrane, dissociation appeared to track the timescale of temperature elevation and image acquisition in our apparatus (approximately 2 minutes), while re-association in some cells was significantly slower. It would be useful to investigate factors such as diffusion and potential low-affinity contacts between unpartnered TlpA strands, their homodimeric partners and other constituents of the cell as potential determinants of these kinetics. With further optimization, thermomers promise to provide a high degree of control over a wide range of cellular processes with the versatile application of thermal energy.

## ACKNOWLEDGEMENTS

The authors thank Mohamad Abedi and Andres Collazo for helpful discussions. Microscopy was performed at the Biological Imaging Facility of the Beckman Institute at Caltech. This research was supported by the Defense Advanced Research Projects Agency (D14AP0050), the Sontag Foundation and the Army Institute for Collaborative Biotechnologies (W911NF-19-D-0001). D.I.P. was supported by the NIH fellowship for Predoctoral Training in Biology and Chemistry (T32GM007616).

## SUPPLEMENTARY MATERIALS

Online Supplementary Materials Include Methods, Supplementary Figures 1-13 and Supplementary Table 1.

## REFERENCES

(1) Piraner, D. I.; Farhadi, A.; Davis, H. C.; Wu, D.; Maresca, D.; Szablowski, J. O.; Shapiro, M. G. Going Deeper: Biomolecular Tools for Acoustic and Magnetic Imaging and Control of Cellular Function. Biochemistry 2017, 56 (39). https://doi.org/10.1021/acs.biochem.7b00443.

(2) Berggård, T.; Linse, S.; James, P. Methods for the Detection and Analysis of Protein – Protein Interactions. 2007, 2833–2842. https://doi.org/10.1002/pmic.200700131.

(3) Spencer, D. M.; Wandless, T. J.; Schreiber, S. L.; Crabtree, G. R. Controlling Signal Transduction with Synthetic Ligands. Science (80-.). 1993, 262 (5136), 1019–1024.

(4) Muthuswamy, S. K.; Gilman, M.; Brugge, J. S. Controlled Dimerization of ErbB Receptors Provides Evidence for Differential Signaling by Homo- and Heterodimers. Mol. Cell. Biol. 1999, 19 (10), 6845–6857.

(5) Wu, C.-Y.; Roybal, K. T.; Puchner, E. M.; Onuffer, J.; Lim, W. A. Remote Control of Therapeutic T Cells through a Small Molecule-Gated Chimeric Receptor. Science (80-.). 2015, 350 (6258). https://doi.org/10.1126/science.aab4077.

(6) Moser, B. A.; Esser-Kahn, A. P. A Photoactivatable Innate Immune Receptor for Optogenetic Inflammation. ACS Chem. Biol. 2017, 12 (2), 347–350. https://doi.org/10.1021/acschembio.6b01012.

(7) Bugaj, L. J.; Choksi, A. T.; Mesuda, C. K.; Kane, R. S.; Schaffer, D. V. Optogenetic Protein Clustering and Signaling Activation in Mammalian Cells. 2013, 10 (3). https://doi.org/10.1038/nmeth.2360.

(8) Maresca, D.; Lakshmanan, A.; Abedi, M.; Bar-Zion, A.; Farhadi, A.; Lu, G. J.; Szablowski, J. O.; Wu, D.; Yoo, S.; Shapiro, M. G. Biomolecular Ultrasound and Sonogenetics. Annu. Rev. Chem. Biomol. Eng. 2018, 9, 229–252. https://doi.org/10.1146/annurev-chembioeng-060817-084034.

(9) Guilhon, E.; Voisin, P.; de Zwart, J. a; Quesson, B.; Salomir, R.; Maurange, C.; Bouchaud, V.; Smirnov, P.; de Verneuil, H.; Vekris, a; et al. Spatial and Temporal Control of Transgene Expression in Vivo Using a Heat-Sensitive Promoter and MRI-Guided Focused Ultrasound. J. Gene Med. 2003, 5 (4), 333–342. https://doi.org/10.1002/jgm.345.

(10) Liu, R. Y.; Corry, P. M.; Lee, Y. J. Regulation of Chemical Stress-Induced Hsp70 Gene Expression in Murine L929 Cells. J. Cell Sci. 1994, 107, 2209–2214.

(11) Krajewski, S. S.; Narberhaus, F. Temperature-Driven Differential Gene Expression by RNA Thermosensors. Biochim. Biophys. Acta - Gene Regul. Mech. 2014, 1839 (10), 978–988. https://doi.org/10.1016/j.bbagrm.2014.03.006.

(12) Piraner, D. I.; Abedi, M. H.; Moser, B. A.; Lee-Gosselin, A.; Shapiro, M. G. Tunable Thermal Bioswitches for in Vivo Control of Microbial Therapeutics. Nat. Chem. Biol. 2017, 13 (1), 75–80. https://doi.org/10.1038/nchembio.2233.

(13) Royal, D. C.; Bianchi, L.; Royal, M. A.; Lizzio, M.; Mukherjee, G.; Nunez, Y. O.; Driscoll, M. Temperature-Sensitive Mutant of the Caenorhabditis Elegans Neurotoxic MEC-4(d) DEG/ENaC Channel Identifies a Site Required for Trafficking or Surface Maintenance. J. Biol. Chem. 2005, 280 (51), 41976–41986. https://doi.org/10.1074/jbc.M510732200.

(14) Cox, V. T.; Baylies, M. K. Specification of Individual Slouch Muscle Progenitors in Drosophila Requires Sequential Wingless Signaling. Development 2005, 132, 713–724. https://doi.org/10.1242/dev.01610.

(15) Hurme, R.; Berndt, K. D.; Normark, S. J.; Rhen, M. A Proteinaceous Gene Regulatory Thermometer in Salmonella. Cell 1997, 90 (1), 55–64.

(16) Stanton, B. Z.; Chory, E. J.; Crabtree, G. R. Chemically Induced Proximity in Biology and Medicine. Science (80-.). 2018, 359 (6380), 1–9. https://doi.org/10.1126/science.aao5902.

(17) Mason, J. M.; Arndt, K. M. Coiled Coil Domains: Stability, Specificity, and Biological Implications. Chembiochem 2004, 5 (2), 170–176. https://doi.org/10.1002/cbic.200300781.

(18) Koski, P.; Saarilahti, H.; Sukupolvi, S.; Taira, S.; Riikonen, P.; Osterlund, K.; Hurme, R.; Rhen, M. A New Alpha-Helical Coiled Coil Protein Encoded by the Salmonella Typhimurium Virulence Plasmid. J. Biol. Chem. 1992, 267 (17), 12258–12265.

(19) Lupas, a; Van Dyke, M.; Stock, J. Predicting Coiled Coils from Protein Sequences. Science (80-.). 1991, 252 (5009), 1162–1164. https://doi.org/10.1126/science.252.5009.1162.

(20) Tripet, B.; Yu, L.; Bautista, D. L.; Wong, W. Y.; Irvin. R. T.; Hodges, R. S. Engineering a de Novo-Designed Coiled-Coil Heterodimerization Domain off the Rapid Detection, Purification and Characterization of Recombinantly Expressed Peptides and Proteins. Protein Eng. 1996, 9 (11), 1029–1042. https://doi.org/10.1093/protein/9.11.1029.

(21) Azuma, Y.; Ku, T.; Yasunaga, J.; Imanishi, M.; Tanaka, G.; Nakase, I.; Maruno, T.; Kobayashi, Y.; Arndt, K. M.; Matsuoka, M.; et al. Controlling Leucine-Zipper Partner Recognition in Cells through Modification of a–g Interactions. Chem Commun 2014, 50 (48), 6364–6367. https://doi.org/10.1039/c4cc00555d.

(22) Kawe, M.; Horn, U.; Plückthun, A. Facile Promoter Deletion in Escherichia Coli in Response to Leaky Expression of Very Robust and Benign Proteins from Common Expression Vectors. Microb. Cell Fact. 2009, 8 (8), 1–8. https://doi.org/10.1186/1475-2859-8-8.

(23) Silva, F.; Queiroz, J. A.; Domingues, F. C. Evaluating Metabolic Stress and Plasmid Stability in Plasmid DNA Production by Escherichia Coli. Biotechnol. Adv. 2012, 30 (3), 691–708. https://doi.org/10.1016/j.biotechadv.2011.12.005.

(24) Hurme, R.; Namork, E.; Nurmiaho-Lassila, E.-L.; Rhen, M. Intermediate Filament-like Network Formed in Vitro by a Bacterial Coiled Coil Protein. J. Biol. Chem. 1994, 269 (14), 10675–10682.

(25) Delviks, K. A.; Pathak, V. K. Effect of Distance between Homologous Sequences and 3’ Homology on the Frequency of Retroviral Reverse Transcriptase Template Switching. J. Virol. 1999, 73 (10), 7923–7932.

(26) DeRose, R.; Miyamoto, T.; Inoue, T. Manipulating Signaling at Will: Chemically-Inducible Dimerization (CID) Techniques Resolve Problems in Cell Biology. Pflügers Arch. - Eur. J. Physiol. 2013, 465 (3), 409–417. https://doi.org/10.1007/s00424-012-1208-6.

(27) Wu, C.-Y.; Rupp, L. J.; Roybal, K. T.; Lim, W. A. Synthetic Biology Approaches to Engineer T Cells. Curr. Opin. Immunol. 2015, 35, 123–130. https://doi.org/10.1016/j.coi.2015.06.015.

(28) Gilbert, C.; Ellis, T. Biological Engineered Living Materials – Growing Functional Materials with Genetically-Programmable Properties. ACS Synth. Biol. 2019, 8 (1), 1–15. https://doi.org/10.1021/acssynbio.8b00423.

(29) Apostolovic, B.; Danial, M.; Klok, H.-A. Coiled Coils: Attractive Protein Folding Motifs for the Fabrication of Self-Assembled, Responsive and Bioactive Materials. Chem. Soc. Rev. 2010, 39, 3541–3575. https://doi.org/10.1039/b914339b.

(30) Muller, K. M.; Arndt, K. M.; Alber, T. Protein Fusions to Coiled-Coil Domains. In Methods in Enzymology; 2000; Vol. 328, pp 261–282. https://doi.org/10.1016/S0076-6879(00)28402-4.

(31) Yuzawa, S.; Mizuno, T.; Tanaka, T. Activating an Enzyme by an Engineered Coiled Coil Switch. Chem. Eur. J. 2006, 12, 7345–7352. https://doi.org/10.1002/chem.200600007.

(32) Woolley, G. A.; Jaikaran, A. S. I.; Berezovski, M.; Calarco, J. P.; Krylov, S. N.; Smart, O. S.; Kumita, J. R. Reversible Photocontrol of DNA Binding by a Designed GCN4-BZIP Protein. Biochemistry 2006, 45, 6075–6084.

