## Supplementary Information for "Modular Thermal Control of Protein Dimerization"

**Table of Contents**
Methods
Supplementary Figures S1-S13
Supplementary Table 1
Supplementary References

### *Methods*

*Plasmid Construction and Molecular Biology*
All plasmids were designed using SnapGene (GSL Biotech) and assembled via KLD mutagenesis or Gibson Assembly using enzymes from New England Biolabs. All plasmids and sequences will be deposited to Addgene. After assembly, constructs were transformed into NEB Turbo and NEB Stable *E. coli* (New England Biolabs) for growth and plasmid preparation. Constructs containing long homologous regions such, including all plasmids containing two TlpA ORFs, were propagated using NEB Stable. Thermal gene expression assays were performed in NEB Stable *E. coli*. Bacterial reporters of gene expression referred to in the text as GFP and RFP are mWasabi[1] and mCherry[2], respectively. The mammalian fusion protein fluorophore referred to in the text and figures as RFP is mScarlet-I[3]. TlpA, mCherry and mWasabi were obtained from our previous work[4]. mScarlet-I was obtained from Addgene (pmScarlet-i_C1, plasmid #85044). Coiled-coil structure prediction and helical wheel diagram annotation was performed using the software DrawCoil 1.0[5]. In the creation of dual-expression GFP/RFP thermal reporter plasmids such as that described in Fig. 1e, an additional terminator was placed upstream of each pTlpA promoter to suppress crosstalk from the weak upstream transcription previously observed from this element[4]. nhTlpA was designed using a homemade script to minimize codon homology while retaining protein sequence identity. Subsequently, 11 nucleotides were altered manually to minimize short repeats that prevented custom gene synthesis. The nhTlpA$_{39}$ gBlock was synthesized by Integrated DNA Technologies.

*Bacterial Thermal Regulation Assay*
Determination of temperature-dependent gene expression was performed as described previously[4]. Briefly, saturated precultures were diluted to $OD_{600}$ = 0.1 and propagated at 30 °C until reaching $OD_{600}$ = 0.3 as measured via Nanodrop 2000c (Thermo Scientific), at which point 25 uL aliquots were dispensed into PCR tubes with transparent caps (Bio-Rad) and incubated for 12 hours in a thermal gradient using a Bio-Rad C1000 Touch thermocycler. After thermal stimulus, fluorescence

was measured using a Stratagene MX3005p qPCR (Agilent), after which cultures were diluted 4x, transferred into microplates (Costar black / clear bottom), and measured for $OD_{600}$ using a Molecular Devices SpectraMax M5 plate reader. The background-corrected F/OD is reported as described previously[4].

*Protein Expression and Purification for CD Spectroscopy*

pET26b-based expression constructs were transformed into BL21-DE3 *E. coli* and grown on kanamycin-selective plates. Saturated overnight cultures were diluted 1 mL into 400 mL expression cultures and induced with a final IPTG (Sigma Aldrich) concentration of 1 mM at $OD_{600}$ = 0.6. After 24 hours of expression at 25 °C, cultures were harvested by centrifugation using a JLA-16.250 rotor (Beckman Coulter) at 6,000 rpm and 4 °C for 8 minutes. Pellets were lysed using the detergent Solulyse in Tris Buffer (Genlantis) and debris was pelleted by centrifugation at 35,343 rcf in a JS-24.38 rotor (Beckman Coulter). Polyhistidine-tagged proteins were purified on an AKTA purifier (GE Healthcare) using 1 mL cOmplete columns (Roche) and buffer exchanged into 1x PBS (Corning) using Zeba spin desalting columns. Concentration was determined using the Pierce 660nm Protein Assay (Thermo Fisher Scientific) and proteins were stored at 4 °C until use. Proteins were subsequently analyzed within 24 hours of purification.

*Circular Dichroism Spectroscopy*

CD melting curves were taken using an Aviv Circular Dichroism Spectrophotometer (Model 60DS) at 222 nm with 0.1 minute equilibration time and 5 second averaging time. Purified proteins were diluted to 3 μM in 1x PBS and measured in a 1 mm quartz cuvette.

*Temperature-dependent protein fluorescence measurement*

Fluorescent proteins were purified as described above and diluted to 1 μM for analysis. 25 μL samples were placed in N=3 replicates in PCR strips with optically transparent caps (Bio Rad) into a Stratagene MX3005p qPCR (Agilent) for intensity mesaurements. Filter sets used for red, green, and blue proteins were ROX, FAM, and ATTO, respectively. Sample fluorescence was measured continuously as temperature was ramped from 25 °C to 50 °C in 1 °C increments and with 1 minute of equilibration time at each increment.

*Mass-based validation of heterodimerization*

Dual TlpA expression constructs were transformed into BL21-DE3 cells (NEB) and grown as 1 mL precultures in 2xYT/ampicillin for 20-24 hours at 30 °C in an Infors Multitron with shaking at 250 rpm. 10 μL saturated cultures were diluted into 4 mL 2xYT/ampicillin and returned to 30 °C. At $OD_{600}$ – 0.6 to 0.7, cultures were induced with 4 μL of 1 M IPTG (Sigma Aldrich) and returned to 30 °C for 12 hours, at which point they were transferred into 2 mL centrifuge tubes (USA Scientific) and pelleted in a Beckman Microfuge 20 at maximum speed for 1 minute. Supernatant was carefully and completely aspirated with a pipette, and the pellet was weighed and frozen at -20 °C for at least 20 minutes. After thawing, Solulyse in Tris Buffer (Genlantis) was added at 10 μL per 1 mg. Pellets were gently resuspended via pipetting and shaken in an Eppendorf ThermoMixer at room temperature (800 rpm for 20 minutes). Subsequently the tubes were spun at 13,000 rcf for 10 minutes and the lysate was diluted 5-fold in Solulyse in Tris Buffer. A pilot Western blot was performed and total TlpA band intensity was quantified for each variant, after which loading amounts for all variants were normalized to that of the wild type via dilution in Solulyse. For crosslinking, 1 μL of 50 mM $CuCl_2$ (Sigma Aldrich) was added to 10 μL lysate in an Eppendorf microcentrifuge tube. The lysate and $CuCl_2$ solution were pre-heated separately for 5 minutes prior to co-incubation. Subsequently, the lysate and crosslinker mixture was shaken at 800 rpm for 10 minutes in an Eppendorf ThermoMixer at the desired temperature. After 10 minutes of $CuCl_2$-catalyzed crosslinking, the reaction was quenched with 11 μL Laemmli buffer (Bio Rad). For uncrosslinked samples, 10 uL Laemmli buffer was added to the lysate at the appropriate temperature. SDS-PAGEwas performed using 7.5% pre-cast polyacrylamide gels (Bio Rad) run at 75 V for 140 minutes. Western blotting was performed using the Transblot Turbo apparatus and nitrocellulose membrane kit (Bio Rad). Transfer was performed at 25 V for 7 minutes. Membranes were blocked with 5% w/v Blotto milk (Santa Cruz Biotechnology) in 0.05% TBS-Tween for 1 hour at room temperature. Primary staining was performed using the mouse anti-HA sc-7392 antibody (Santa Cruz Biotech) overnight at 4 °C. Blots were then washed three times for 15 minutes at 4 °C with 0.05% TBS-Tween and stained for 4 hours with goat anti-mouse IgG-HRP sc-2005 (Santa Cruz Biotech) at room temperature. After three 15-minute washes, HRP visualization was performed using Supersignal West Pico PLUS reagent (Thermo Fisher Scientific). Imaging was performed in a Bio-Rad ChemiDoc MP gel imager.

*Mammalian cell culture*
K562 cells (gift of D. Baltimore) were cultured in RPMI 1640 media (Thermo Fisher Scientific) with 1x Penicillin/Streptomycin (Corning). Transient transfection was performed using Lonza 4D nucleofection with SF Cell Line buffer and the pre-programmed K562 protocol. Lentivirus was prepared using a third-generation viral vector and helper plasmids (gifts of D. Baltimore). Virus was packaged in HEK293T cells grown in 10 cm dishes after 2 days of transfection and concentrated via the Lenti-X reagent (Takara Bio). Infection was performed by resuspending viral pellets in 250 uL RPMI and spinfecting 1E6 K562 cells in 1 mL RPMI with 10 μL virus at 800 xg, 30 °C, for 90 minutes. Experiments were performed at least five passages after infection.

*Live cell microscopy*
Delta-T dishes (Bioptechs) were coated with 400 μL 0.1 mg/mL Poly-D Lysine (Sigma Aldrich) for 30 minutes. Meanwhile, 1E6 K562 cells were pelleted at 300 rcf for 5 minutes and resuspended in staining solution (1x PBS or HBSS with 2.5 μg/mL Hoechst 33342, Thermo Fisher Scientific). For Golgi staining, the solution (in HBSS) also contained BODIPY FL $C_5$-Ceramide complexed to BSA (Thermo Fisher Scientific, 50 nM final concentration). Cells were stained at room temperature for 10 minutes before being pelleted and resuspended in 1 mL RPMI 1640. For co-localization experiments, the staining solution (in HBSS) also contained CellBrite Fix 488 at 1x concentration as described in the product manual, and staining was performed at 37 °C for 10 minutes. After at least 20 minutes of coating, PDL was aspirated from the Delta-T dishes, which were then rinsed once with 1x PBS and dried. Cells were then transferred to the Delta-T dishes, which where adhesively affixed to the swinging plates of an Allegra X-12 centrifuge with SX4750 rotor (Beckman Coulter) and centrifugated at 30 °C for 15 minutes at 300 rcf. Imaging was performed at the California Institute of Technology confocal microscopy facility using an LSM880 (Zeiss) with Airyscan. Cells with sufficient overall brightness to discern membrane contrast were imaged; the membrane localization of dimmer cells could not be discerned in our thermal imaging configuration but was observable under higher magnification on a conventional glass slide (**Supplementary Fig. S12**). Delta-T dishes were mounted onto the thermal stage interfaced with a Bioptechs Delta T4 Culture Dish Controller and imaged using a 1080-378 C-Achroplan 40x/0.80 W objective. Airyscan processing was performed in 2D mode using default settings.

*Image analysis*

Image analysis was performed using the Zeiss Zen Black software for pre-processing and CellProfiler[6] for correlation quantification. Images were manually cropped to include only a single cell per ROI, with approximately 408x408 pixel FOVs. For cells with poor attachment to the plate, resulting in position offsets between the red, blue, and green channels, frame alignment between the red and green channels was performed in Zen Black. All available transformations were sampled and the best-aligned transformation on a per-cell basis was used. Blue channel alignment was performed in CellProfiler on a per-cell basis for images with obvious displacement of the nucleus using the Mutual Information module, correlating the blue channel image with the inverted red channel image. Exported images were then loaded in CellProfiler and analyzed using a custom pipeline (**Supplementary File F1)**. Briefly, cell boundaries were determined from red channel using Hoescht 33342-stained nuclei as primary seed objects. Colocalization was calculated for the ROI defined between the outer cell membrane and the nucleus. The nucleus was excluded because it acts as a diffusion barrier to TlpA-RFP but not to free mScarlet-I. For the free mScarlet-I cell line in **Fig. 4f**, a modified pipeline using the CellBrite Fix 488 stain for cell boundary determination was used to improve the detection of cell edges (**Supplementary File F2**). For the Direct Membrane Fusion data set in **Fig. 4f**, the 44 °C point was acquired in a separate experiment using the same cell line and is consistent with the 43 °C and 45 °C data points of the original data set (**Supplementary Fig. S13**).

### *Supplementary Figures S1-S14*

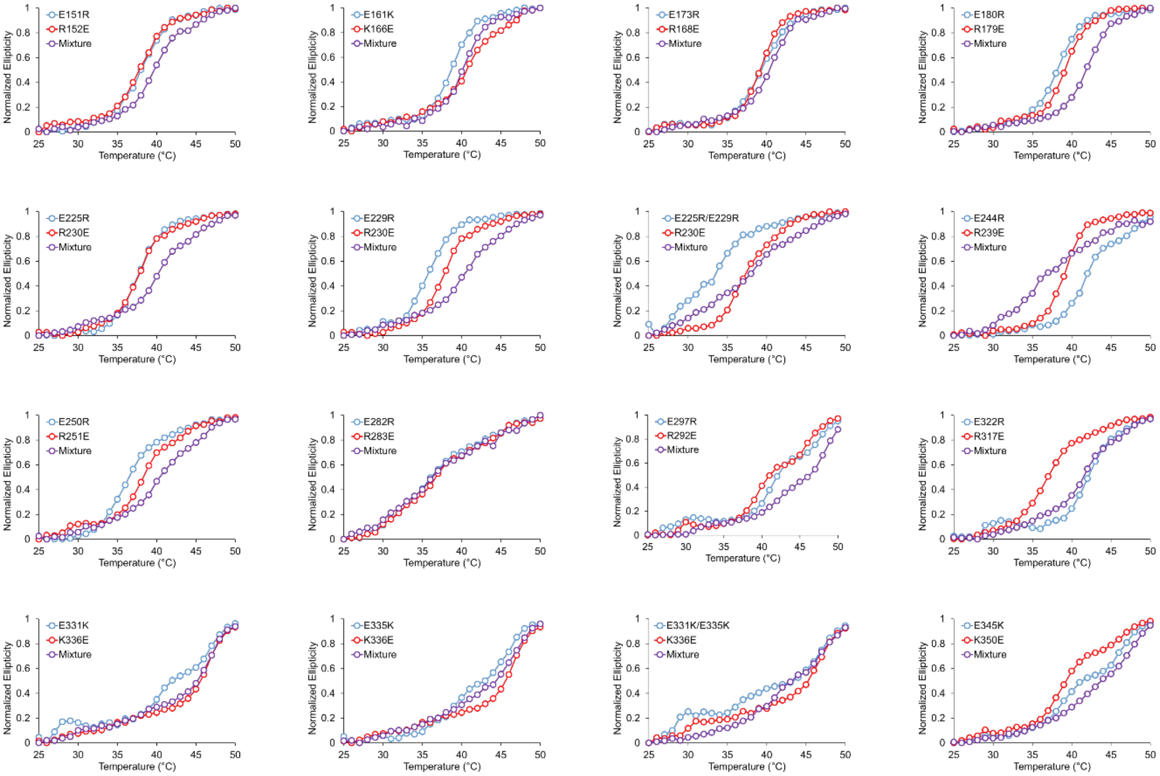


**Supplementary Figure S1**: Circular dichroism melting curves of engineered TlpA variants. Coiled-coil fragments corresponding to residues 69-359 of TlpA were purified from *E. coli* and assayed via CD spectroscopy. Monitoring the ellipticity at 222 nm, corresponding the prototypical peak of the α-helical spectrum, enables tracking the conformation of TlpA as it transitions from the dimeric coiled-coil state to a monomeric random coil configuration.

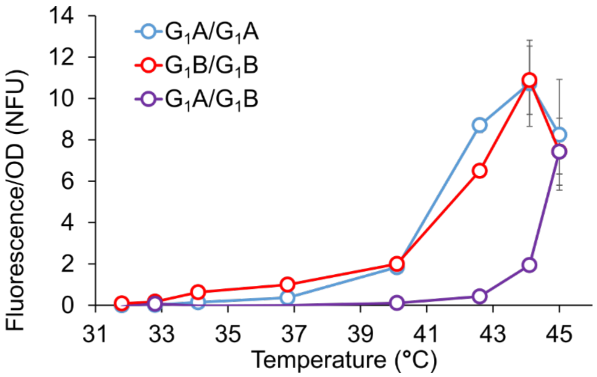


**Supplementary Figure S2**: Thermal RFP expression profiles of the constructs shown in **Fig. 2b** (n = 3). Error bars represent ± s.e.m. Where not seen they are smaller than the symbol.

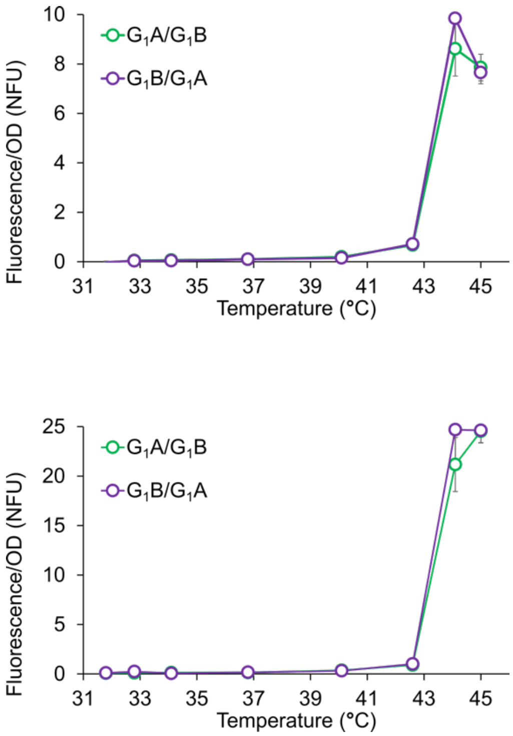


**Supplementary Figure S3**: Positional independence of the TlpA co-expression construct. The positions of TlpA-$G_1$A and TlpA-$G_1$B within the heterodimer co-expression construct in **Fig. 2b** were exchanged and the thermal GFP (top) and RFP (bottom) expression profile was determined (n = 4). No significant differences were observed in the gene expression profile, thereby excluding position-dependent effects on TlpA behavior within the circuit. Error bars represent ± s.e.m. Where not seen they are smaller than the symbol.

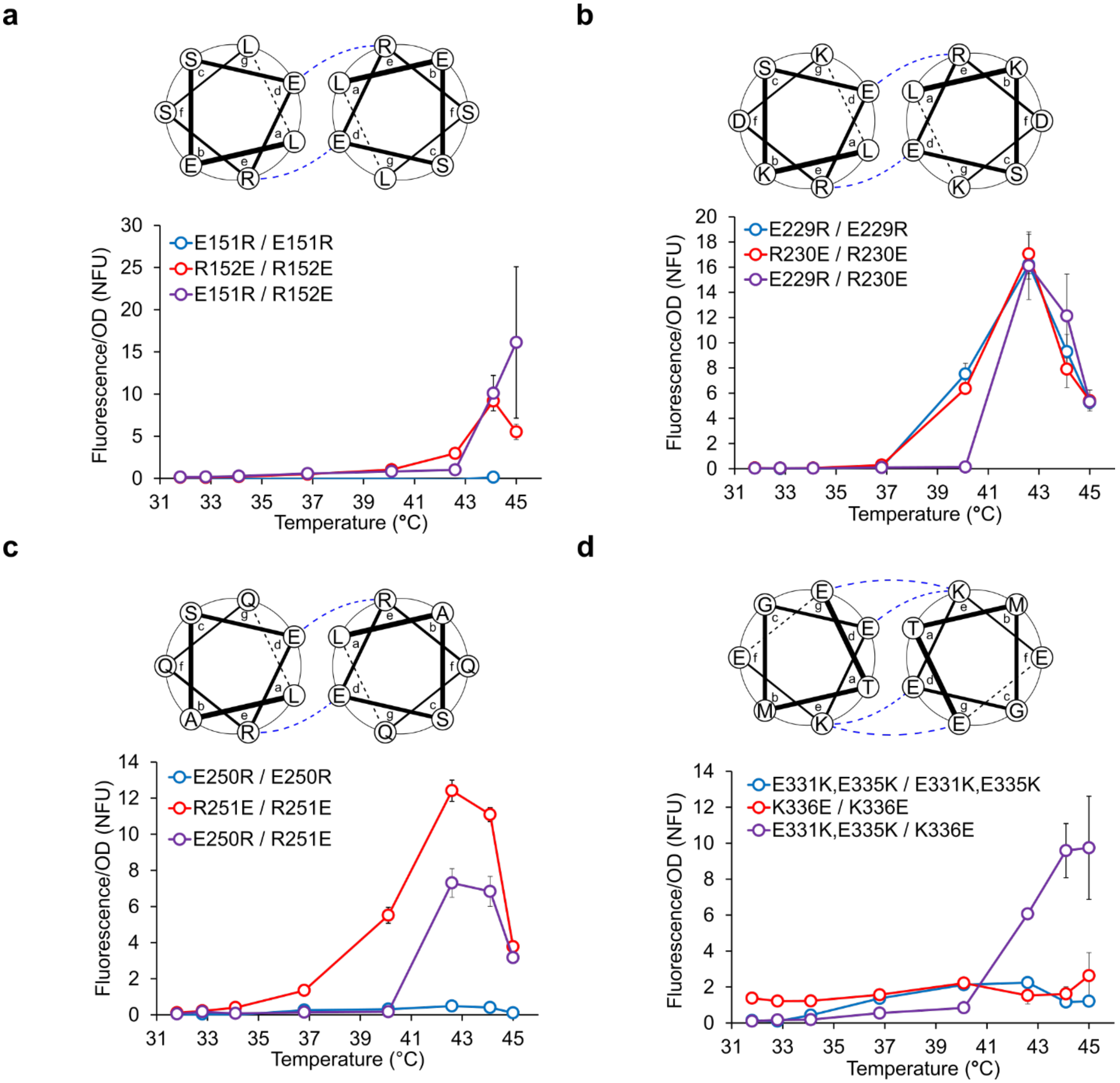


**Supplementary Figure S4:** Screening of rationally designed mutant panel via bacterial thermal gene expression assay. Four positions in the TlpA coiled-coil were selected for mutagenesis based on the predicted similarity of their ionic interaction pattern to the $G_1A/G_1B$ mutant pair according to the heptad repeat register prediction of Koski *et al*. The following mutations were examined: **a)** E151R and R152E **b)** E229R and R230E, **c)** E250R and R251E, and **d)** E331K/E335K and K336E. The predicted interaction pattern of the wild type protein is depicted (top), and the thermal GFP expression profile is reported (bottom). N = 3. The E229R/R230E and E250R/R251E pairs were selected for introduction into the TlpA-$G_1A$ and $G_1B$ variants. Error bars represent ± s.e.m. Where not seen they are smaller than the symbol.

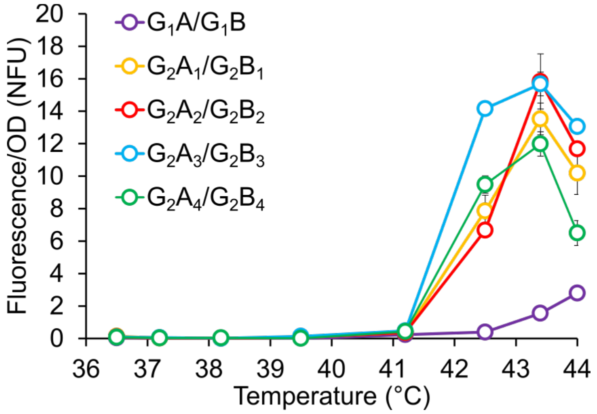


**Supplementary Figure S5**: Thermal RFP expression profiles of the first and second-generation heterodimers shown in **Fig. 2c** (n = 3). Only the construct containing one copy of each heterodimeric strands are depicted because the $X_nA/X_nB$ homodimeric construct were unable to propagate without accumulating deletion mutations in the TlpA promoters or fluorescent protein open reading frames. Error bars represent ± s.e.m. Where not seen they are smaller than the symbol.

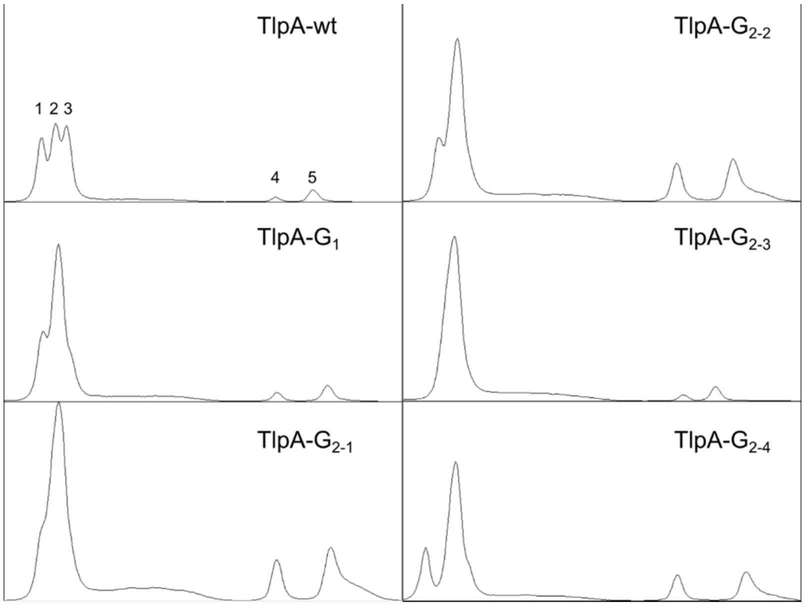


**Supplementary Figure S6**: Western blot band intensity profiles of co-expressed full and truncated TlpA strands in **Fig. 3c**. Note that samples lacking a distinct band corresponding to the truncated homodimer (3), such as TlpA-$G_1$, nevertheless display higher band intensity for the truncated uncrosslinked strand (5) relative to the full-length uncrosslinked species (4), confirming that the lack of a low molecular weight homodimer at position 3 results from reduced homodimer affinity rather than full depletion of the light TlpA strand by heterodimer pairing. This is consistent with cationic and anionic TlpA variants having different homodimerization affinities at 37 °C.

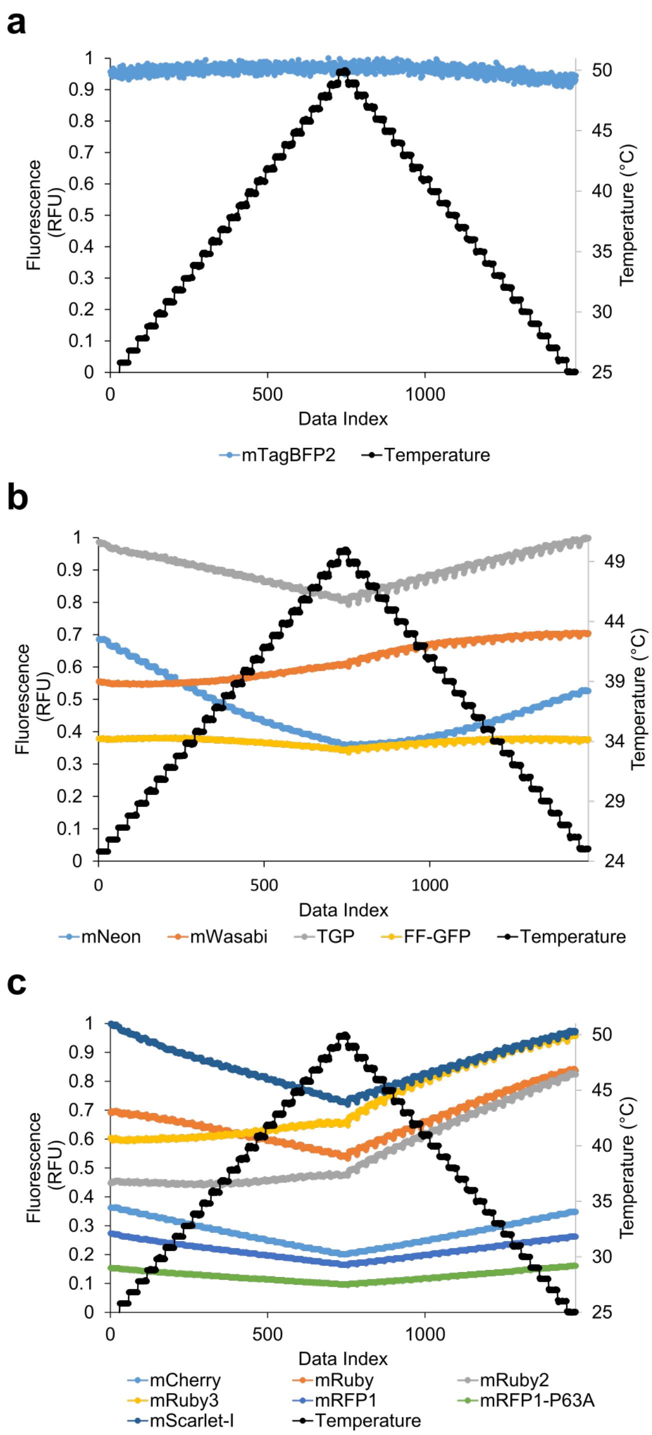


**Supplementary Figure S7**: Thermal stability of a panel of blue (**a**), green (**b**), and red (**c**) fluorescent proteins. Proteins were prepared in equimolar concentrations and their fluorescence was measured in an rtPCR thermocycler upon a thermal ramp from 25 °C to 50 °C and subsequent re-annealing to 25 °C, with readings taken continuously over 1 minute intervals. Signal intensity is normalized to the maximum for each given experiment. Because different filter sets were used for the three classes of proteins, relative brightness does not correlate between the red, green, and blue channels. While some proteins such as FF-GFP demonstrated more stable signal over the temperature range tested, the overall brightness was maximal in mScarlet-I and TGP. However, TGP demonstrated significant aggregation when expressed as an untagged cytosolic protein in mammalian cells so mScarlet-I was chosen as the reporter for subsequent experiments.

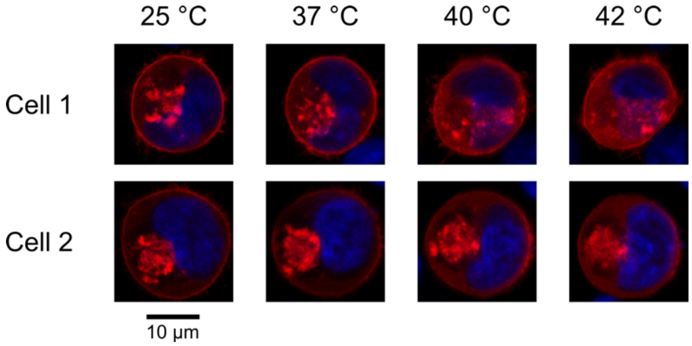

**Supplementary Figure S8**: Additional replicates for TlpA membrane localization experiment in **Figure 4b**. Pixel intensity was normalized to the maximal per-image value.

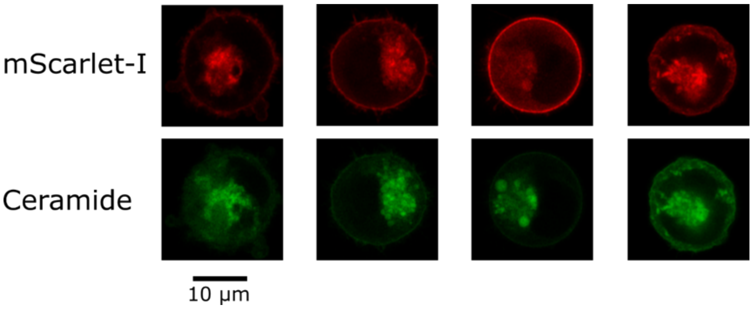

**Supplementary Figure S9**: K562 cells transfected with the construct shown in **Fig. 4a** were stained with BODIPY-C5-Ceramide to label the Golgi transport pathway. Staining morphology was similar to the localization of the mScarlet-I TlpA cargo protein. Four representative cells are shown. Pixel intensity was normalized to the maximal per-image value.

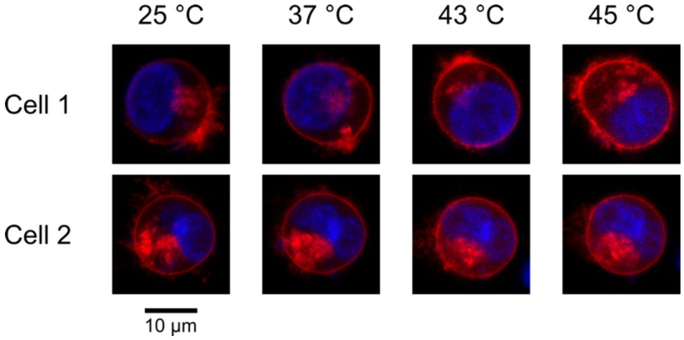

**Supplementary Figure S10**: Additional replicates of K562 cells transfected with a construct bearing directly palmitoylated mScarlet-I, as in **Fig. 4d**. Robust membrane localization was observed up to 45 °C in most cells. Data presented are representative of two separate experiments. Pixel intensity was normalized to the maximal per-image value.

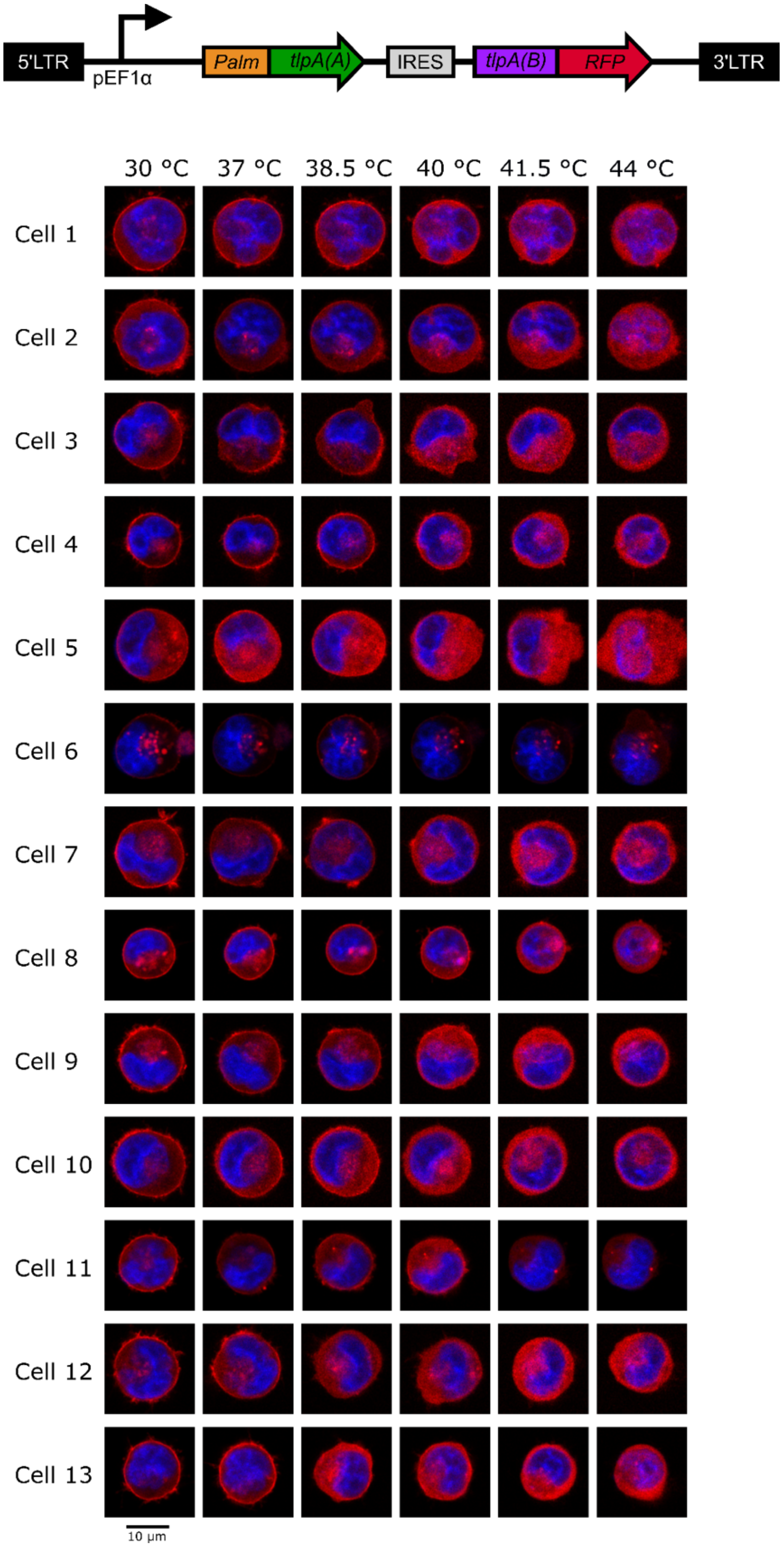


**Supplementary Figure S11**: Top: Lentiviral construct for membrane-localized RFP delivery containing nonhomologous $TlpA_{39}$-$G_2A_3$. Bottom: Fluorescence images at different temperatures of K562 cells lentivirally transduced with the nonhomologous TlpA-mediated membrane localization system. Pixel intensity was normalized to the maximal per-image value.

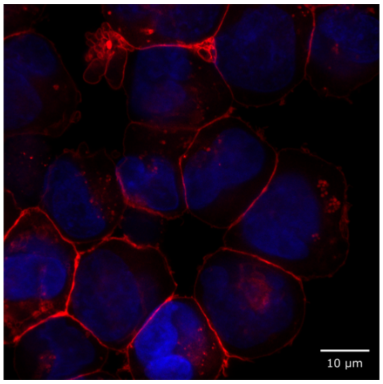


**Supplementary Figure S12**: Confocal imaging of K562 cell line transduced with the lentiviral construct depicted in **Supplementary Figure S11**. Cells were pelleted, deposited on a glass slide, sealed with a cover slip, and imaged on an LSM880 with a Plan-Apochromat 63x/1.4 Oil DIC M27 objective with oil immersion. Note that all cells, regardless of local cytoplasmic or membrane brightness, display visible RFP accumulation along the plasma membrane.

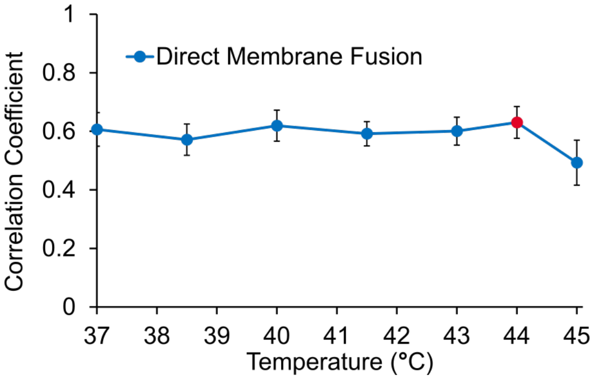


**Supplementary Figure S13**: CellProfiler quantification of the two data sets contributing to the Direct Membrane Fusion curve in **Fig. 4f**. The data point acquired separately at 44 °C is highlighted in red. Error bars represent ± s.e.m.

**Supplementary Table 1**: Second-generation TlpA mutants utilized in this study.

| Mutant | Mutations |
|---|---|
| $G_2A_1$ | E180R + E229R |
| $G_2B_1$ | R179E + R230E |
| $G_2A_2$ | E180R + R230E |
| $G_2B_2$ | R179E + E229R |
| $G_2A_3$ | E180R + E250R |
| $G_2B_3$ | R179E + R251E |
| $G_2A_4$ | E180R + R251E |
| $G_2B_4$ | R179E + E250R |

***Supplementary References***


(1) Ai, H.; Olenych, S. G.; Wong, P.; Davidson, M. W.; Campbell, R. E. Hue-Shifted Monomeric Variants of Clavularia Cyan Fluorescent Protein: Identification of the Molecular Determinants of Color and Applications in Fluorescence Imaging. *BMC Biol.* **2008**, *6*, 13. https://doi.org/10.1186/1741-7007-6-13.

(2) Shaner, N. C.; Campbell, R. E.; Steinbach, P. A.; Giepmans, B. N. G.; Palmer, A. E.; Tsien, R. Y. Improved Monomeric Red, Orange and Yellow Fluorescent Proteins Derived from Discosoma Sp. Red Fluorescent Protein. *Nat. Biotechnol.* **2004**, *22* (12), 1567–1572. https://doi.org/10.1038/nbt1037.

(3) Mastop, M.; Bindels, D. S.; Shaner, N. C.; Postma, M.; Gadella, T. W. J.; Goedhart, J. Characterization of a Spectrally Diverse Set of Fluorescent Proteins as FRET Acceptors for MTurquoise2. *Sci. Rep.* **2017**, *7* (1), 11999. https://doi.org/10.1038/s41598-017-12212-x.

(4) Piraner, D. I.; Abedi, M. H.; Moser, B. A.; Lee-Gosselin, A.; Shapiro, M. G. Tunable Thermal Bioswitches for in Vivo Control of Microbial Therapeutics. *Nat. Chem. Biol.* **2017**, *13* (1), 75–80. https://doi.org/10.1038/nchembio.2233.

(5) Grigoryan, G.; Keating, A. Structural Specificity in Coiled-Coil Interactions. *Curr. Opin. Struct. Biol.* **2008**, *18* (4), 477–483. https://doi.org/10.1016/j.sbi.2008.04.008.Structural.

(6) Carpenter, A. E.; Jones, T. R.; Lamprecht, M. R.; Clarke, C.; Kang, I. H.; Friman, O.; Guertin, D. A.; Chang, J. H.; Lindquist, R. A.; Moffat, J.; et al. CellProfiler: Image Analysis Software for Identifying and Quantifying Cell Phenotypes. *Genome Biol.* **2006**, *7* (10). https://doi.org/10.1186/gb-2006-7-10-r100.
